# Cytotoxic and mutagenic capacity of TTO and terpinen-4-ol in oral squamous cell carcinoma

**DOI:** 10.1101/2020.01.03.893735

**Authors:** Nicole Casalle, Cleverton Roberto de Andrade

## Abstract

The essential oil of *Melaleuca Alternifolia* (tea tree oil - TTO) consists of about 100 components, and the highest concentration are terpinen-4-ol. Studies of their cytotoxic capacity have shown effect on malignant neoplastic lineages. The aim of this study was to evaluate the cytotoxic and mutagenic capacity of TTO and main soluble components, terpinen-4-ol and gama-terpinene in cell cultures. Two lineages of oral squamous cell carcinoma and a keratinocyte cell were analyzed: (1) colorimetric analysis Metiltetrazolium (MTT); (2) Micronucleus assay. The results were expressed as susceptibility tests and degree of mutagenicity. The statistical test used in the analysis was *one-way* ANOVA (Tukey test). The IC_50_ values obtained from the MTT analysis of cells exposed to TTO were 0.2% for HaCaT, 0.14% for HSC-3, and 0.17% for SCC-25. For exposure to terpinen-4ol, IC_50_ values were 0.5%, 0.3% and 0.45% for HaCaT, HSC-3 and SCC-25, respectively. The gamma-terpinene didn’t show significant cytotoxic activity, therefore it was impossible to calculate the IC_50_. TTO and terpinen-4-ol was unable to produce mutagenicity in all the lineages. In conclusion, both the TTO and terpinen-4-ol had cytotoxic capacity on HaCaT, HSC-3 and SCC-25. TTO and terpinen-4-ol wasn’t mutagenic. In this sense, our study provides new perspectives on the potential use of TTO and terpinen-4-ol for the development of new alternative therapies to treat topically locally oral squamous cell carcinoma. This study can be related

## Introduction

Oral cancer has been growing day by day and currently occupies 11th worldwide position between the most common carcinomas^1^. Squamous cell carcinoma are responsable for over 90% of oral cancers. Treatment for this cancer may envolve highly mutilating surgeries, chemotherapy and radiotherapy, which are procedures with high cost and low accessibility. As a result, studies try to identify new options for treatment that can improve the survival of patients with oral squamous cell carcinoma (SCS) ^2, 3^. In this context, experiments with plant-derived components have been performed to identify new drugs of potential use as auxiliaries to the proposed treatment ^4, 5^. In fact, new chemotherapy drugs originally extracted from plants continue to enter the market and are being analyzed for their usefulness against different cancers, including squamous cell carcinoma.

*Melaleuca alternifolia* is a herbal medicine that stands out in different studies for medicinal purposes ^6^. *Melaleuca* essential oil (tea tree oil-TTO) is composed of a complex mixture of approximately 100 components ^7^, of which the highest concentration components are terpinen-4-ol (39-48%), gamma-terpinene (17-18%) and alpha-terpinene (8-13%). The oil is considered non-toxic, with a pleasant odor and is widely used in healing or anti-infectious products. It is extracted from the plant by hydro distillation or steam distillation ^6^. Its lipophilic capacity and consequent skin penetration suggest the possibility of topical use ^8, 9^. The monoterpenes, delivered from TTO, have also been studied for their anticâncer activity. Some of them have shown promising results in the treatment and prevention of skin câncer and leukemia, and others types of cancer such as pancreas, colon and breast in rodents ^10^.

This study evaluated the cytotoxicity of TTO and its major soluble componentes: terpinen-4-ol and gamma-terpinene, against head and neck squamous cell carcinoma (SCC) by possible alteration of mitochondrial activity (oxidative respiration, bromine [3-(4,5-dimethylthiazol-2yl) -2,5-diphenyl-2H tetrazolate assay - MTT). MTT assay is based on the effect of compound / extract on glycid cell metabolism through mitochondrial dehydrogenases. The mitochondrial viability and consequently cell viability is quantitated buy cell’s ability to convert MTT salt (water-soluble and yellow) in formazan (purple precipitate and water-insoluble) via mitochondrial dehydrogenases^11^.

Subsequently, it was evaluated if the cytotoxic effect causes nuclear damage, at chromosome level, these chromosomal mutations were analyzed by the Micronucleus test (MN). Eventually it is possible that damage to the DNA of the cell doesn’t lead to apoptosis or necrosis. These damages can cause chromosomal mutations, which are essential factors to carcinogenesis ^12^, therefore, we consider it important to identify the possible mutagenic capacity of our test substances in concentrations lower than those capable of cell death. The micronucleus test (MN) has emerged as one of the confirmatory method for assessing chromosomal damage. MNs are expressed during the cellular division and are identified in binucleated phase, when chromosomal damage occurs, fragments or whole chromosomes doesn’t migrate to the spindle poles during mitosis. A special method was developed to identify these cells in binucleated phase, which occurs when there is a blockage of cytokinesis by Cytochalasin-B (CIT-B). CIT-B is an actin polymerization inhibitor of cytoplasmic division by blocking the formation of contractile microfilaments and cell cleavage into two daughter cells – cytokinesis, without affecting nuclear division. The use of CIT-B results in an accumulation of binucleate cells from cells that have passed by only one division cycle. Having the formation of MNs, they will also be contained in the cytoplasm, are identical to the nucleus, but morphologically smaller. Events that are related to the formation of micronucleus can cause activation of proto-oncogenes or deletion of tumor suppressor genes or DNA repair, resulting in the malignant transformation. MNs frequency represents an index composed for chromosomal aberrations in a given cell population, Schmid et al. ^13^ (2007), proposed in 1975 the use as a short-term test to evaluate the genotoxic potential of chemical and physical agents ^14^.

## Materials and methods

### Tea tree oil and components

The composition of the TTO (Sigma, MO, USA) was analyzed by gas chromatography for oil quality control and to obtain the percentages of its main components. The identification of the relative concentrations of the components was used to calculate the initial concentration to perform the experiments with the two components of higher concentration (terpinen-4-ol and gamma-terpinene).

The ISO 4730 values of the oil components and the composition obtained in the chromatography analysis are expressed in Fig.1. The percentage values for terpinen-4-ol and gamma-terpinene were, respectively: 47.66% and 23.58%, based on the chromatography performed.

**Fig. 1.**
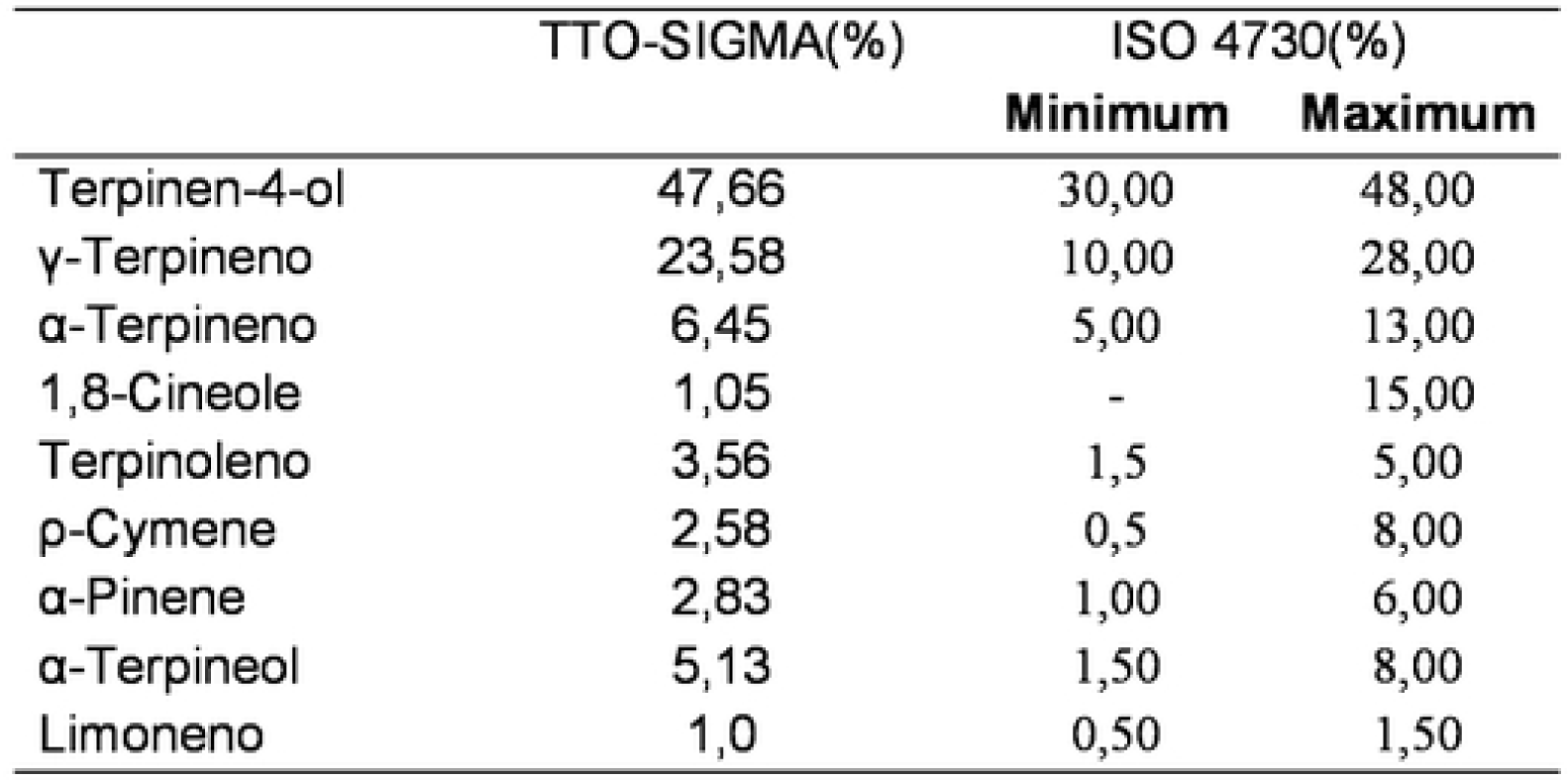
Tea tree oil composition obtained in the chromatography. Data from oil chromatography, in comparison with the ISO 4730 values.

### Cell lines

Two head and neck squamous cell carcinoma lines, SCC-25 (Squamous cell carcinoma) and HSC-3 (Human oral squamous cell carcinoma) and an epitelial human keratinocytes Cell line (Human Keratinocyte Cell line - HaCaT) were provided by the Department of Oral Diagnosis of the Piracicaba School of Dentistry (UNICAMP). HaCaT was maintained in DMEM/F10 with 10% fetal bovine serum (SFB) and 1% penicillin-streptomycin and frozen at -80 ° C with 10% dimethyl sulfoxide (DMSO). SCC-25 line was maintained in DMEM/F12 with 10% fetal bovine serum (FBS) and penicillin-streptomycin 1%, 400ng of hydrocortisone per ml and frozen at -80°C with 10% dimethylsulfoxide (DMSO). Finally, HSC-3 line was maintained in DMEM/F12 with 10% FBS, 1% penicillin-streptomycin, 400ng of hydrocortisone per ml, 0.01g of Vitamin C for each 200 ml of medium and frozen at -80°C with 10% DMSO. The following experiments were performed in triplicate.

### MTT cell viability assay

This analysis served as a parameter for producing the IC_50_, which was used as the standard concentration for subsequent experiments ^15^. Thus, this first analysis had more dilutions of TTO and their soluble portions. This experiment also served to determine the inclusion of soluble portions, the component that could not calculate the IC_50_ was not included in the next experiment. In addition, as TTO is the initial viability analysis product, none of the component was analyzed at a higher concentration than that found in TTO (proportional relative concentration tested in TTO).

Cells were seeded in 96-well plates (1×10^5^ cells/wells) in complete medium. On the following day, different concentrations of TTO and soluble portions were added to the cells in triplicates. It was exposed to TTO at decreasing concentrations (1/2) from the initial concentration of 2% (1%, 0.5%, 0.25%, 0.125%, 0.062%, 0.031%) for 24 hours, in all cases, combining 0.4% dimethyl sulfoxide diluent (DMSO) into the culture medium. For the main soluble portions, we use the chromatography result to calculate the relative concentrations of each product and adjust for the experiments. In the case of terpinen-4-ol the initial concentration used was 1% (0.5%, 0.25%, 0.125%, 0.062%, 0.031%, 0.015%), for the same time as exposure to TTO and for gamma-terpinene was 0.5% (0.25%, 0.125%, 0.062%, 0.031%, 0.015%). For the soluble portions, no diluent was used in the culture medium as it was water soluble. At 24 h later, the medium was replaced by fresh media (100 μL per well) containing 0.5 mg/ml 3-(4,5-dimethylthiazol-2-yl)-2,5-diphenyltetrazolium bromide (MTT) and incubated for 4h at 37°C. MTT-formazan crystals were dissolved by the addition of 100 μl of DMSO. Absorbance was recorded with a spectrophotometer (BIO-RAD, model 3550-UV, microplate reader Hercules, California, USA) for 570 nm wavelength plates. After this analysis, the data were tabulated using the negative control considered as 100% viability. Subsequently, the results were transformed to logarithmic basis and the IC_50_ was calculated (50% of viable cells), identifying the concentration required to obtain 50% of viable cells in TTO and soluble portions. This data was used for the MN experiment,

### Micronucleus test

Cells were seeded in 96-well plates (4×10^3^ cells/wells) in complete medium. On the following day, different concentrations of TTO and soluble portions were added to the cells in triplicates. The concentrations were based on MTT analysis. Three concentrations below the IC_50_ of each linage were used. The experiment was performed with 24 hours exposure, and the concentrations tested in each linage were IC_20_, IC_10_, IC_05_. The positive control was 0.1 mM Hydrogen Peroxide (PC). Citocalasin B was then placed for 36 hours in the incubator at 37°C. After this time, the plate was centrifuged for 5 minutes at 4,000 rpm. Citocalasin B was then removed and each well was washed with 1X PBS (400µL / well). The cells were fixed with absolute ethanol (100µL / well) for 30 minutes at room temperature. 100µl / well of FITC (1µg / mL) was added over 30 minutes at room temperature. The wells were washed again with 1X PBS (400µl / well) and placed 50µl / well Hoescht (10µg / mL) for 15 minutes. The images were immediately acquired and analyzed using the IN Cell Analyzer 2000 equipment (GE Healthcare), located at the Proteomic Nucleus of the Araraquara Faculty of Pharmaceutical Sciences - UNESP.

### Statistics

T-test and chi-square analysis were performed comparing the results obtained by *Melaleuca alternifolia* oil, soluble portion and controls. For comparation, the results was applied to the analysis of variance one-way ANOVA with Tukey post test. Analysis were performed using a statistical program, Graphpad PrismTM6.0 (La Jolla, California, USA). The mean values were obtained by the arithmetic sum of the individual results divided by the number of repetitions. However, statistical analysis were performed with the pure numbers obtained in each individual experiment. Student’s t-test was used with p <0.05 for significance (* p <0.05 ** p <0.01 *** p <0.001).

### Study approval

This study was approved by the Ethics Committee on Human Research (CEP), by Araraquara School of Dentistry - UNESP.

## Results

### TTO, terpinen-4-ol and gama-terpineno as an anti-cancer agents

TTO and terpinen-4-ol were able to reduce cell viability in all lines (Fig.2). Increased cytotoxicity was observed at higher concentrations, while at lower concentrations, the percentages of cell death were low. There was no significant variation between them. HaCaT line demonstrated cell viability reduction by more than 50% of cells until a concentration of 0.25% when exposed to TTO (IC50: 0.2) (Fig.2-A) and 0.5% when exposed to terpinen-4-ol, (IC 50: 0.5) (Fig. 2-B). For HSC-3, the inhibitory concentration was approximately 0.125% when exposed to TTO (IC50: 0.14) (Fig.2-D) and 0.25% when exposed to terpinen-4-ol (IC50: 0, 3) (Fig.2-E). SCC-25 line showed cell viability reduction by more than 50% until a concentration of 0.25% when exposed to TTO (IC50: 0.17) (Fig.2-G) and 0.5 % when exposed to terpinen-4-ol (IC50: 0.45) (Fig.2-H). For terpinen-4ol IC_50_ concentrations, were twice as high as for TTO in all lineages. Gamma-terpinene was not able to provide reduction of cell viability of oral carcinoma cells, so it was not possible to calculate IC_50_ for gamma-terpinene. (Fig.2-C.F.I).

**Fig 2.**
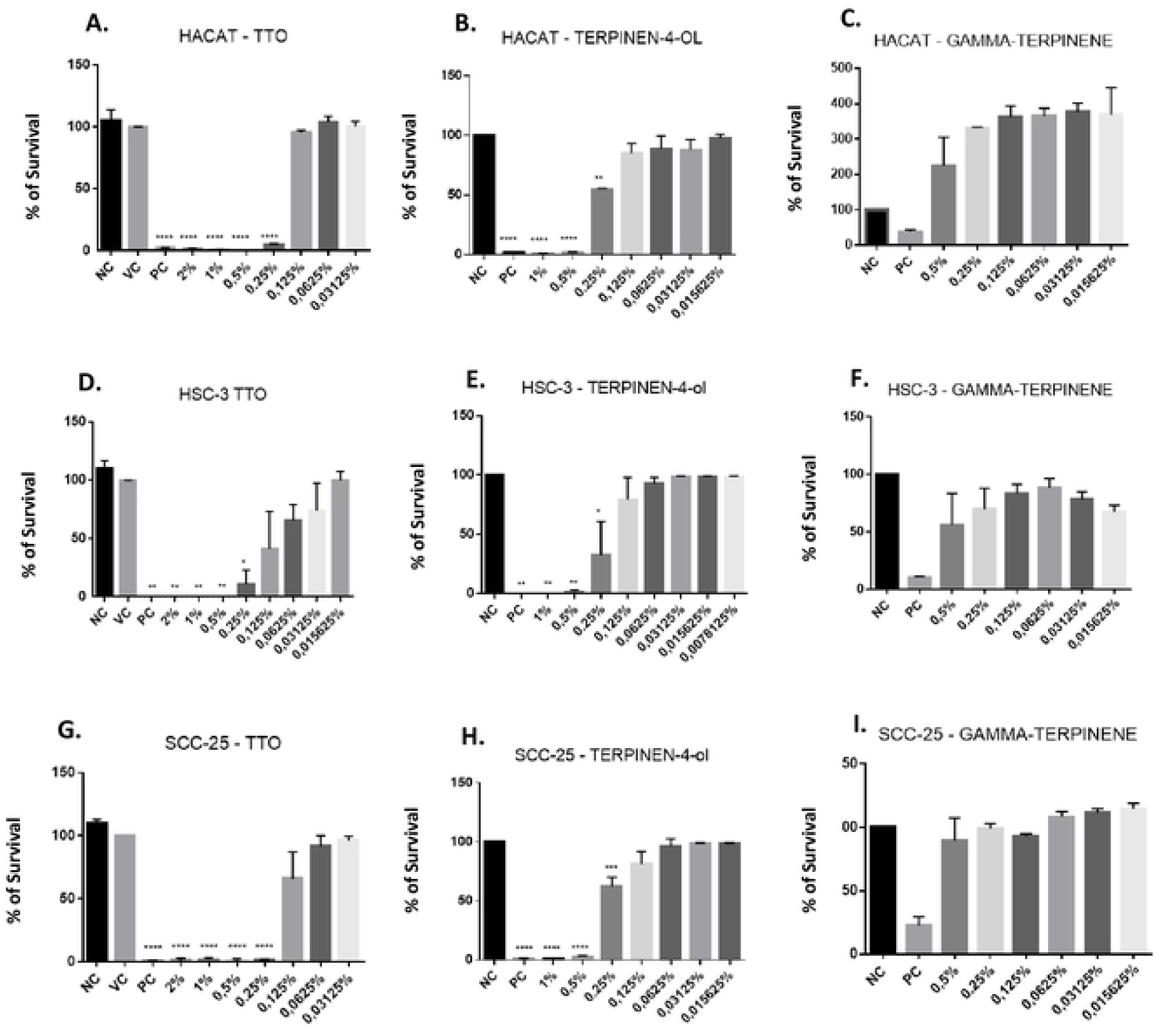
MTT assay. Data refer to the means of three independent experiments and standard error (M ± EP). NC, negative control (DMEM), VC, vehicle control (DMEM + 0.4% DMSO), PC, positive control (Doxorubicin). TTO and terpinen-4-ol were tested at different concentrations for 24 hours in each lineage. Statistical analysis was performed using One-way ANOVA with Tukey’s posttest (Treated vs NC/ VC). %, p <0.05, ** p <0.01, *** p <0.001, **** p <0.000.

### TTO and terpinen-4-ol mutagenic capacity

TTO and terpinen-4-ol were not mutagenic (Fig.3). IC_20_, IC_10_ and IC_05_ were statistically different from PC, and the three concentrations tested showed no statistical difference among themselves or against the NC and VC groups, showing that the concentrations were not mutagenic against any of the analyzed lines.

**Fig. 3.**
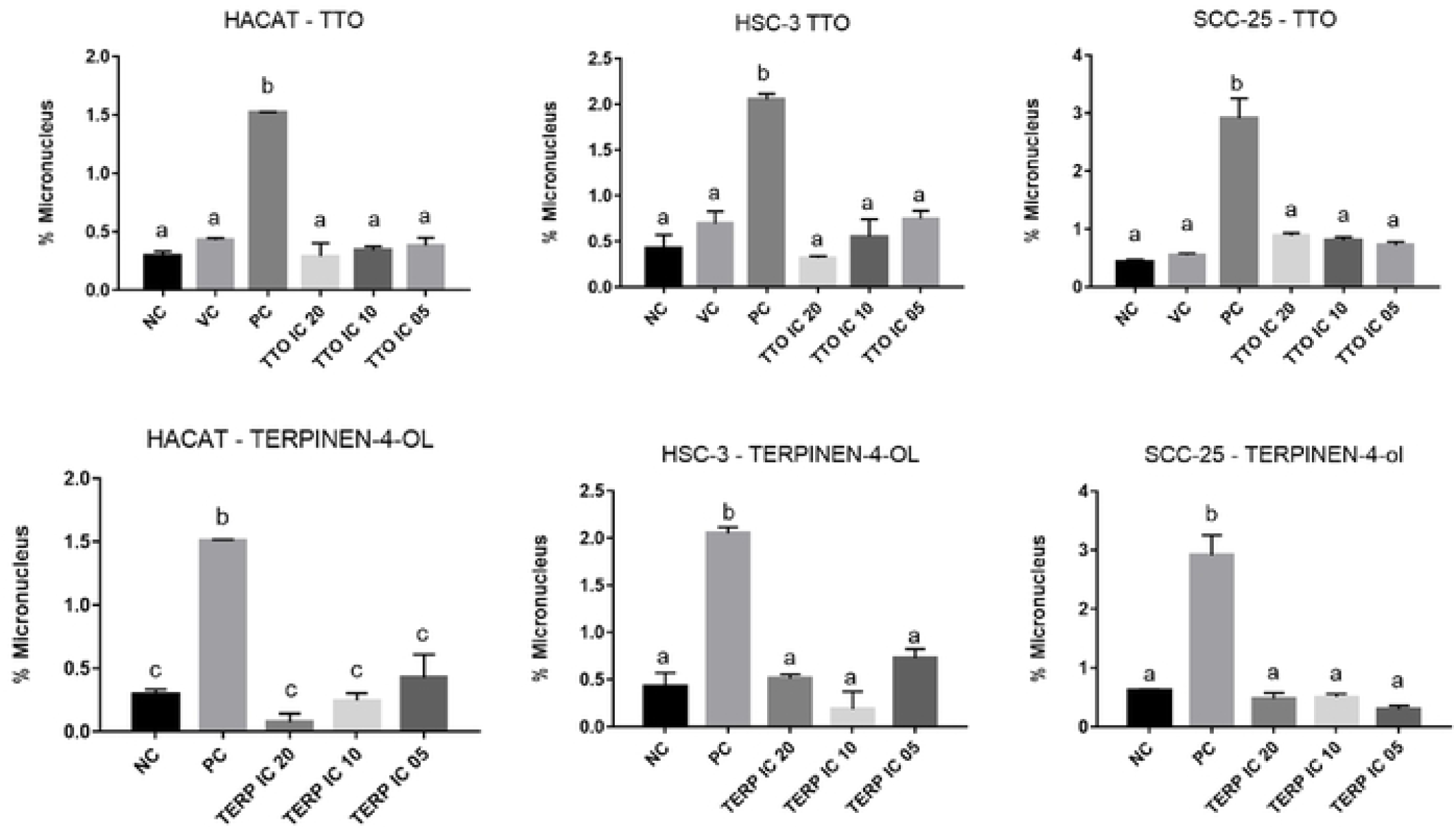
Micronucleus test. All MN test data refer to the averages of three independent experiments (M ± EP) and were expressed as the percentage of micronucleus founded in each group tested. NC, negative control (DMEM), VC, vehicle control (DMEM + 0.4% DMSO), PC, positive control (0.1mM Hydrogen Peroxide, for 15 minutes exposure). The images were immediately acquired and analyzed using the IN Cell Analyzer 2000 equipment (GE Healthcare), located at the Proteomic Nucleus of the Araraquara Faculty of Pharmaceutical Sciences - UNESP. Statistical analyzes were performed with Graphpad PrismTM6.0 software (La Jolla, California, and were performed by One-way ANOVA with Tukey post-test). %, * a <0.05, ** b <0.01, *** c <0.001, ****.

## Discussion

In recent decades, much has been said about the use of medicines from natural extracts^6^. The alarming rise in cancer cases around the world, and the limitations of current treatments, have made increased the search for treatments with natural medicines that have antitumor action.

Doll-Boscardin et al. ^16^ (2012), confirmed in an in vitro study, the cytotoxic and antitumor potential of *Eucalyptus benthamii* essential oil and some related terpenes (alpha-terpinen, terpinen-4-ol and gamma-terpinene) in three different tumor lineages. The results indicated that the cytotoxic effect caused cell death by apoptosis. In addition, there was inhibition of cell proliferation in consequence of cytotoxicity.

Krifa et al. ^17^ (2015), studied the essential oil (EO) extracted from the *Pituranthos Torturou*s plant and main components, sabinene, alpha-pinene, limonene and terpinen-4ol, on melanoma lineages. The results showed that EO was able to inhibit tumor growth associated with apoptotic features, including nuclear condensation, suggesting a potential drug for cancer treatment.

Although the essential oil of *Melaleuca alternifolia* (tea tree oil - TTO) and their soluble portions, are widely used for their known bactericidal action, antifungal and anti-inflammatory, there’s no much information about its anti-tumor capacity. Some studies have been conducted to prove this ability, but there are no reports in the literature about the antitumor effect of TTO and soluble portions in squamous cell carcinoma cell lines.

In the present study, experiments were performed to evaluate cytotoxicity and degree of mutagenicity in the squamous cell carcinoma lines, HSC-3 and SCC-25 compared to the HaCaT line.

The cytotoxic assay used for our study was colorimetric analysis of Metiltetrazolium (MTT). Based on this assay, we found that both TTO and terpinen-4-ol were able to reduce viability in all strains. The initial TTO concentration used was 2%, followed by seven dilutions (1/2). While for terpinen-4-ol, the initial concentration was half, 1%, according to its relative percentage found by gas chromatography.

Increased cytotoxicity was observed at higher concentrations, while at lower concentrations, cell death percentages were low, suggesting a dose-dependent effect for both TTO and terpinen-4-ol.

The IC_50_ found for HaCaT line was 0.2%. This value was higher than those found for squamous cell carcinoma lines, HSC-3 and SCC-25, which were 0.14% and 0.17%, respectively, suggesting greater susceptibility of carcinoma lines. The same occurred after exposure to terpinen-4-ol, where the IC_50_ values of the three lineages doubled, being 0.5% for HaCaT, 0.3% for HSC-3 and 0.45% for SCC-25. This fact occurred because the relative percentage of terpinen-4-ol in the oil is approximately 50%. The gamma-terpinene soluble portion, present in approximately 24% of the TTO, was not able to reduce viability in 50% of the cells, so it was not used in the other analysis.

Differently to the concentrations found in the present study, Hammer et al.^18^ (2006), in a literature review about the cytotoxicity of the essential oil of *Melaleuca alternifolia*, reported that TTO produces toxic effects on human monocyte cells at concentrations ≥ 0.004%. and in human neutrophils ≥ 0.016%. However, there are no data on the exposure time of the compounds used in the study.

Calcabrini et al. ^4^ (2004) also reported using lower TTO concentrations, 0.005% to 0.03%, for MTT analysis in human melanoma cells. The authors reported there was no difference in cell viability at a concentration of 0.005% and until to 0.01%, there was no interference on the growth of human melanoma line tested by them. From 0.02% and 0.03% concentrations was possible to observe large inhibition of cell growth. This study also does not mention the exposure time of TTO and terpinen-4-ol concentrations used.

In study by Greay et al.^5^ (2009), the tested concentrations of TTO, terpinen-4-ol and gamma-terpinene, on mice mesothelioma, melanoma and on fibroblast line, ranged from 0, 01% to 0.15%, exposed to treatments for 24 to 72 hours. The IC_50_ found for TTO was 0.03% and 0.02% for terpinen-4-ol, exposure for 48 hours. These concentrations were less effective against fibroblasts, and the IC_50_ for this lineage was 0.08% for TTO and 0.1% and for terpinen-4-ol in 24 hours exposure. As in the present study, the authors reported that gamma-terpinene and other soluble portions were unable to produce IC_50_. They also mention dose-dependent effect of TTO and terpinen-4-ol, which was also mentioned in the studies by Khaw-on and Banjerdpongchai ^19^ (2012) and Banjerdpongchai and Khaw-on ^20^ (2013).

Previous studies have indicated that terpinen-4-ol is the most active component of *Melaleuca alternifolia* oil. This can be seen in our study, when we proved that terpinen-4-ol was responsible for producing half the IC_50_ found in TTO.

Wu et al. ^21^ (2012), tested several concentrations (0.02% to 0.1%) of terpinen-4-ol in two human cell lines of lung cancer, exposure for 24 hours. The IC_50_ obtained was 0.052% and 0.046%. The authors also mention the dose-dependent effect of terpinen-4-ol.

Recently, Assmann et al. ^22^ (2018) analyzed the cytotoxic and antitumor capacity of TTO and major constituents against breast cancer cells in human and in mice (MCF-7 and 4T1, respectively), as well as cytotoxic action on fibroblasts and mononuclear blood peripheral cells (PBMCs). In this study, the authors also demonstrate the fact that TTO exhibited antitumor effect in vitro against MCF-7 and 4T1 cells, reducing cell viability and modulating apoptotic pathways and stopping the cell cycle of MCF-7 cells.

In order to evaluate possible mutagenic effect of TTO and terpinen-4-ol exposure for 24 hours, we performed the Micronucleus test. We observed that TTO and terpinen-4-ol didn’t cause mutagenicity in any of the tested lineages.

There are no studies about the mutagenic effect of TTO and soluble portions in malignant lines, but the literature review by Hammer et al.^12^ (2006) shows that TTO and its active components indicated low genotoxicity potential in bacterial lines and mammals, a fact that can be evidenced in our study.

## Conclusion

In conclusion, both the TTO and terpinen-4-ol had cytotoxic capacity on HaCaT, HSC-3 and SCC-25. TTO and terpinen-4-ol wasn’t mutagenic. In this sense, our study provides new perspectives on the potential use of TTO and terpinen-4-ol for the development of new alternative therapies to treat topically locally oral squamous cell carcinoma.

## Acknowledgments

The authors thank Dr. Christianne Pienna Soares (Department of Cytology) for providing the Cytology Laboratory of the Faculty of Pharmaceutical Sciences for the work could be done and Dr. Ricardo Della Coletta (Department of Oral Diagnosis) of the Faculty of Dentistry of Piracicaba-UNICAMP for providing the cell lines for our research. This study was supported in part by CNPq.

